# Rod nuclear architecture determines contrast transmission of the retina and behavioral sensitivity in mice

**DOI:** 10.1101/752444

**Authors:** Kaushikaram Subramanian, Martin Weigert, Oliver Borsch, Heike Petzold, Alfonso Garcia, Eugene Myers, Marius Ader, Irina Solovei, Moritz Kreysing

## Abstract

Rod photoreceptors of nocturnal mammals display a striking inversion of nuclear architecture, which has been proposed as an evolutionary adaptation to dark environments. However, the nature of visual benefits and underlying mechanisms remains unclear. It is widely assumed that improvements in nocturnal vision would depend on maximization of photon capture, at the expense of image detail. Here we show that retinal optical quality improves 2-fold during terminal development, which, confirmed by a mouse model, happens due to nuclear inversion.

We further reveal that improved retinal contrast-transmission, rather than photon-budget or resolution, leads to enhanced contrast sensitivity under low light condition. Our findings therefore add functional significance to a prominent exception of nuclear organization and establish retinal contrast-transmission as a decisive determinant of mammalian visual perception.

**One Sentence Summary:** Our study reveals that chromatin compaction in rod cells augments contrast sensitivity in mice.

## Introduction

The structure of the vertebrate retina requires light to pass through multiple cell layers prior to reaching the light-sensitive outer segments of the photoreceptors (Dowling, 1987). In nocturnal mammals, the increased density of rod photoreceptor cells demands a thicker (Němec, Cveková, Burda, Benada, & Peichl, 2007; Peichl, 2005) rod nuclei-containing outer nuclear layer (ONL). For mice, where rods account for around 80% of all retinal cells (Hughes, Enright, Myers, Shen, & Corbo, 2017), this layer of photoreceptor nuclei is 55±5*µ*m thick, thus creating an apparent paradox by acting as a more pronounced barrier for projected images prior to their detection (Fig. 1A). Interestingly, rod nuclei are inverted in nocturnal mammals (Błaszczak, Kreysing, & Guck, 2014; Kreysing, Boyde, Guck, & Chalut, 2010; Solovei et al., 2009; 2013), such that heterochromatin is detached from the nuclear envelope and found in the nuclear center, whereas the less-dense euchromatin is re-located to the nuclear periphery. Given that this nuclear inversion is exclusive to nocturnal mammals and correlates with the light-focusing capabilities of isolated nuclei, it was proposed as an evolutionary adaptation to life under low-light conditions (Błaszczak et al., 2014; Kreysing et al., 2010; Solovei et al., 2009). However, the nature of any visual improvements that could arise from nuclear inversion remains unclear.

**Fig. 1.**
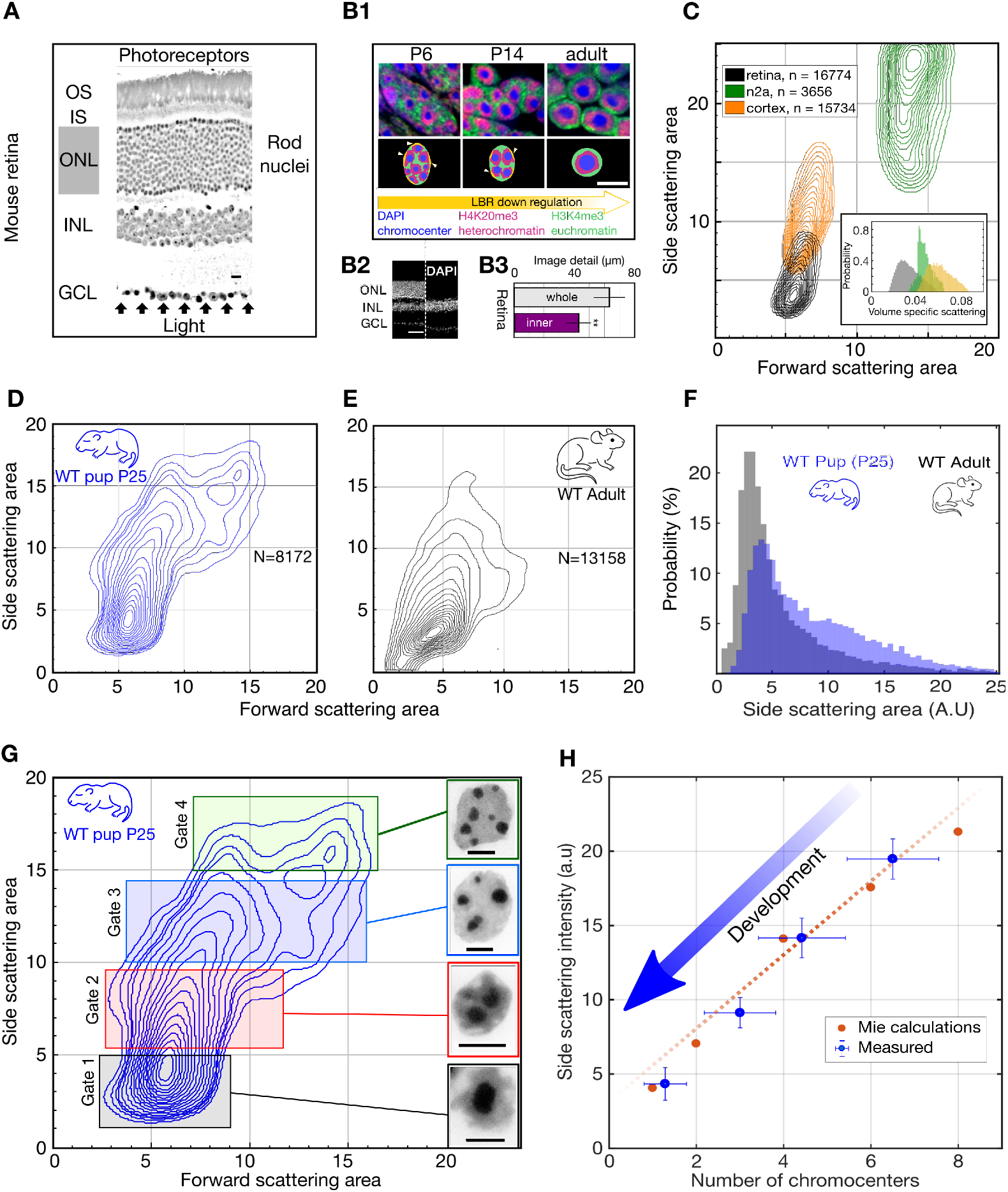
Light scattering by retinal nuclei reduces with chromocenter number during development. ***(A)*** Longitudinal section showing the path of light through the mouse retina, including the rod nuclei dominated outer nuclear layer (ONL). Ganglion cell layer (GCL), inner nuclear layer (INL) and outer nuclear layer (ONL) and the inner and outer segments (IS & OS). **(B1) (top)** Downregulation of the lamina tether LBR (yellow) enables fusion of mobilized chromocenters and thereby an architectural inversion of mouse rod nuclei. **(bottom)** FISH images of rod nuclei stained with DAPI (blue) showing the dense chromocenters, LINE rich heterochromatin (H4K20me3, magenta) and SINE rich euchromatin (H3K4me3, green) **(B2)** DAPI section of WT mouse retina in comparison to a rd1-cpfl1 mouse retina showing the presence of only the inner retina. **(B3)** Quantification of image transmission shows that the inner retina alone (rd1-cpfl1, N=5) *transmits approximately 50% more image* detail than the full retina (N=11), suggesting significant image degradation in the thick outer nuclear layer. **(C)** FACS scattering profiles comparing retinal neurons, cortical neurons and N2a neuroblastoma cells and showing lower light scattering properties of retina neurons. (Inset) Volume specific light scattering is significantly reduced in the retina cell nuclei. **(D, E)** FACS scatter plot for isolated retinal nuclei from WT developmental stage week 3 pup (P25) and adult mice demonstrating stronger large angle scattering by the P25 nuclei. **(F)** Histogram of side scattering in adult and P25 retina depicting a higher side scattering for the developing retinal nuclei. **(G)** Sorting of developmentally maturing nuclei according to different side scattering signal. Insets show representative examples of Hoechst stained nuclei in the corresponding sort fractions. **(H)** Quantification of reduced scattering with chromocenter number is sufficiently explained by a wave optical model of light scattering. (Error bars in **H** show s.d.) Scale bars **A** - 10 µm**. B1, G -** 5 µm, **B2** – 50µm

It is widely assumed that high-sensitivity vision depends on optimized photon capture (Schmucker & Schaeffel, 2004; Warrant & Locket, 2004) and often comes at the expense of image detail (Cronin, Johnsen, Marshall, & Warrant, 2014; Warrant, 1999). Here, we show that nuclear inversion affects a different metric of vision, namely contrast sensitivity under low-light conditions. In particular, we experimentally show that nuclear inversion improves retinal contrast transmission, rather than photon capture or resolution. Advanced optical modelling and large-angle scattering measurements indicate that this enhanced contrast transfer emerges from previously coarse-grained (Błaszczak et al., 2014; Kreysing et al., 2010; Solovei et al., 2009) changes in nuclear granularity, namely a developmental reduction of chromocenter number (Fig. 1B1). Moreover, genetic interventions to change chromocenter number in adult mice reduces contrast transmission through the retina, and compromise nocturnal contrast sensitivity accordingly. Our study therefore adds functional significance to nuclear inversion by establishing retinal contrast transmission as a decisive determinant of mammalian vision.

## Results

### Volume specific light scattering from chromocenters

To test how the presence of densely packed rod nuclei in the light path affects the propagation of light through the retina, we compared transmission of micro-projected stripe images through freshly excised retinae of wild type (WT) (Fig. 1A) and Rd1/Cpfl1 KO mice(Chang et al., 2002), which lack all photoreceptors including the ONL (Fig. 1B2). In the absence of photoreceptors and their nuclei, we observed 49% greater imaged detail (cut-off chosen at 50% residual contrast, Fig. 1B3). Since photoreceptor nuclei contain highly compacted and molecularly dense DNA with significant light-scattering potential (Drezek et al., 2003; Marina, Sanders, & Mourant, 2012; Mourant et al., 2000), while photoreceptor segments have been described as image-preserving waveguides (Enoch, 1961), these findings suggest that light propagation in the mouse retina is significantly impacted, if not dominated, by the highly abundant rod nuclei of the ONL.

We then asked whether retinal cell somata are optically specialized with distinct light scattering properties. A comparison of light scattering by different cell types using high throughput FACS(Feodorova, Koch, Bultman, Michalakis, & Solovei, 2015) measurements revealed that isolated retinal cells scatter substantially less light than neurons of the brain and cultured neuroblastoma cells (Fig. 1C). This trend is seen for forward-scattered light but is even more pronounced for side scattering, which reflects subcellular heterogeneity. Using forward scattering as a measure of cell size indicates that side scattering normalized by volume is also noticeably lower in retinal cells (Fig. 1C, inset). This suggest that retinal cells are indeed optically specialized, as they scatter less light for a given size.

To determine when the low sideward light scattering characteristic of retinal nuclei emerges, we compared the scattering profile of retinal nuclei in P25 pups and adult (12 weeks) mice. We found little or no difference between forward light scattering (Fig. 1D-E), as predicted by earlier models (Błaszczak et al., 2014; Kreysing et al., 2010; Nagelberg et al., 2017). In stark contrast, however, side scattering (measured in a narrow range around 90°), with a strong potential to diminish image contrast, was significantly reduced in adult retinal nuclei compared to the intermediate developmental stage (Fig. 1F). Quantitative analysis of sorted nuclei from P25 retinae further revealed a monotonic relation between chromocenter number and side scattering signal (Fig. 1G). In particular, those nuclei with the lowest number of chromocenters were found to scatter the least. In support of this experimental quantification, a wave-optical Mie model of light-scattering by refractive chromocenters closely reproduced the trend of light scattering reduction with chromocenter fusion (Fig. 1H).

To establish whether rod nuclear inversion is required to cause the developmental reduction in light scattering, we used a transgenic mouse model (TG-LBR) in which heterochromatin remains anchored at the lamina which in turn prevents the complete fusion of chromocenters (Fig. 2A1, A2, 2B, Fig. S2, S3)(Solovei et al., 2013). FACS experiments of nuclei from TG-LBR retinae, in which >70% of the nuclei are successfully arrested (Fig. S4), revealed significantly increased light scattering (Fig. 2C, Fig. 1F). Specifically, the global maximum of the side-scattering was re-located precisely to the position that is characteristic of nuclei isolated from WT pups at P14, which possess a similar number of chromocenters as inversion arrested nuclei (compare Fig. 2B, C, Fig. 1F, G). Because inhibition of chromocenter fusion leads to specific increase in scattering, we conclude that the reduction of light scattering with chromocenter number is causal.

**Fig. 2.**
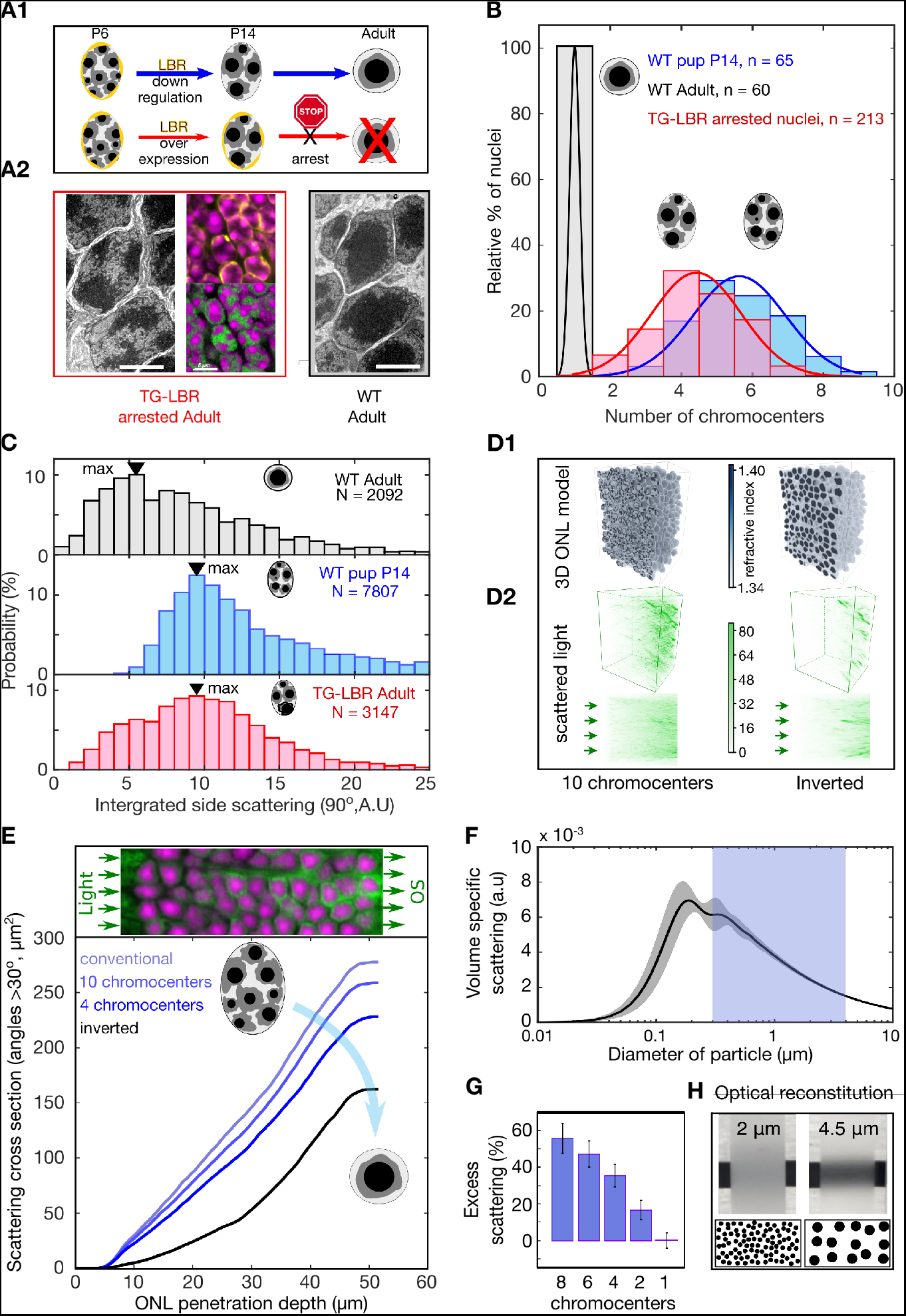
Developmental arrest of chromocenter fusion increases light scattering from rod nuclei in measurements and tissue simulations. (A1) Schematic of the normal rod nuclear WT development and inversion arrested nuclei by LBR overexpression. (A2) EM images illustrating different electron densities in the euchromatic and heterochromatic phase underlying their refractive index (RI) differences (scalebar 5µm). (mid-top) Immunostaining of overexpressed of LBR tethers (yellow), and high-density heterochromatin (DAPI, magenta). (mid-bottom) Heterochromatic chromocenters (DAPI, magenta) and euchromatin (H4K5ac, green) (B) Chromocenter number distribution in LBR overexpressing rod nuclei is drastically different from WT mice, and similar to a developing WT pup (P14). (C) Side scattering assessed by FACS for TG-LBR retina nuclei is higher than that of WT nuclei and comparable to that of a WT P14 nuclei with similar chromocenter numbers. Note the shift of peak value upon LBR overexpression. (D1) 3d RI distribution mapped onto anatomically faithful volumetric ONL images. WT inverted architecture (right, top) and early developmental state (left) (simulation). (D2) (top) Differential simulations of light propagation in the ONL, using same positions and shapes of about 1750 nuclei, but varying chromatin distributions. (bottom) Maximum projection illustrating greater proportions of scattered light (angles >30deg) in the ONL with multiple chromo-centered nuclei. (E) Quantitative analysis of this data. (F) Angle weighted volume specific scattering strength for nuclei models evaluated by Mie scattering theory. (G) Excess scattering occurring in multi-chromocenter nuclei models. (H) Chromocenters scattering reconstituted in an emulsion of silica spheres in glycerol-water mixture. Pictograms reflect accurate number ratio of spheres.

### Improved retinal contrast transmission

Next, we asked how nuclear substructure could affect the optical properties of the ONL. We first approached this via a simulation that built on recent advances in computational optics (Weigert, Subramanian, Bundschuh, Myers, & Kreysing, 2018). This allowed us to specifically change nuclear architecture, while leaving all other parameters, including the morphology and relative positioning of about 1750 two-photon mapped nuclei, unchanged (Fig. 2D1) (Supplementary Methods).

These simulations suggested that especially the sideward scattering (cumulative scattering signal at angles >30 deg) monotonically decreases when 10 chromocenters successfully fuse into one (Fig. 2D2, 2E). Physically, this effect of reduced scattering can be explained by a reduction of volume-specific scattering for weak scatterers in the size regime slightly above one wavelength of light, similar to scattering reduction techniques proposed for transparent sea animals(Johnsen, 2012) (Fig. 2 F, G). Furthermore, a minimal optical ONL model reconstituted from suspended beads of different size but same volume fraction (Supplementary Methods) illustrates how a decreased geometric scattering cross section after fusion leads to reduced scattering-induced veil that helps to prevent contrast losses (Fig. 2H and inset). Taken together these data suggest that nuclear inversion might serve to preserve contrast in retinal transmitted images.

To experimentally quantify the optical quality of the retina with respect to nuclear architecture, we applied the concept of the modulation transfer function (MTF), a standard way to assess image quality of optical instruments (Boreman, 2001). Specifically, MTF indicates how much contrast is maintained in images of increasingly finer sinusoidal stripes (Fig. S5 B, C). We therefore devised an automated optical setup (Fig. S5 A) that allowed us to project video sequences of demagnified sinusoidal stripe patterns through freshly excised retinae and assess the retinal transmitted images for contrast loss. This custom built set-up mimics the optics of the mouse eye, in particular its f-number (Schmucker & Schaeffel, 2004), while circumventing changes of the optical apparatus in-vivo (Fig. S5, Supplementary Materials & Methods).

Strikingly, we found that wildtype retinae improve contrast transmission throughout terminal development, with adult retinae showing consistently elevated MTFs compared to intermediate developmental stages (P14) in which rod nuclei still possess around 5 chromocenters (Fig 3A). In contrast to many lens-based optical systems, retinal MTFs do not display a strict resolution limit. Instead a monotonic decay of retina transmitted contrast indicates scattering induced veil, rather than a frequency cut-off to be the cause of contrast loss (Fig. 3 A, B Fig. S6 A-D). Collected from >1300 high resolution images, this data reveals that, similar to the lens (T. V. Tkatchenko, Shen, & Tkatchenko, 2010), the retina matures towards increasing optical quality during latest developmental stages, with chromocenter fusion as a putative mechanism of veil reduction.

**Fig. 3:**
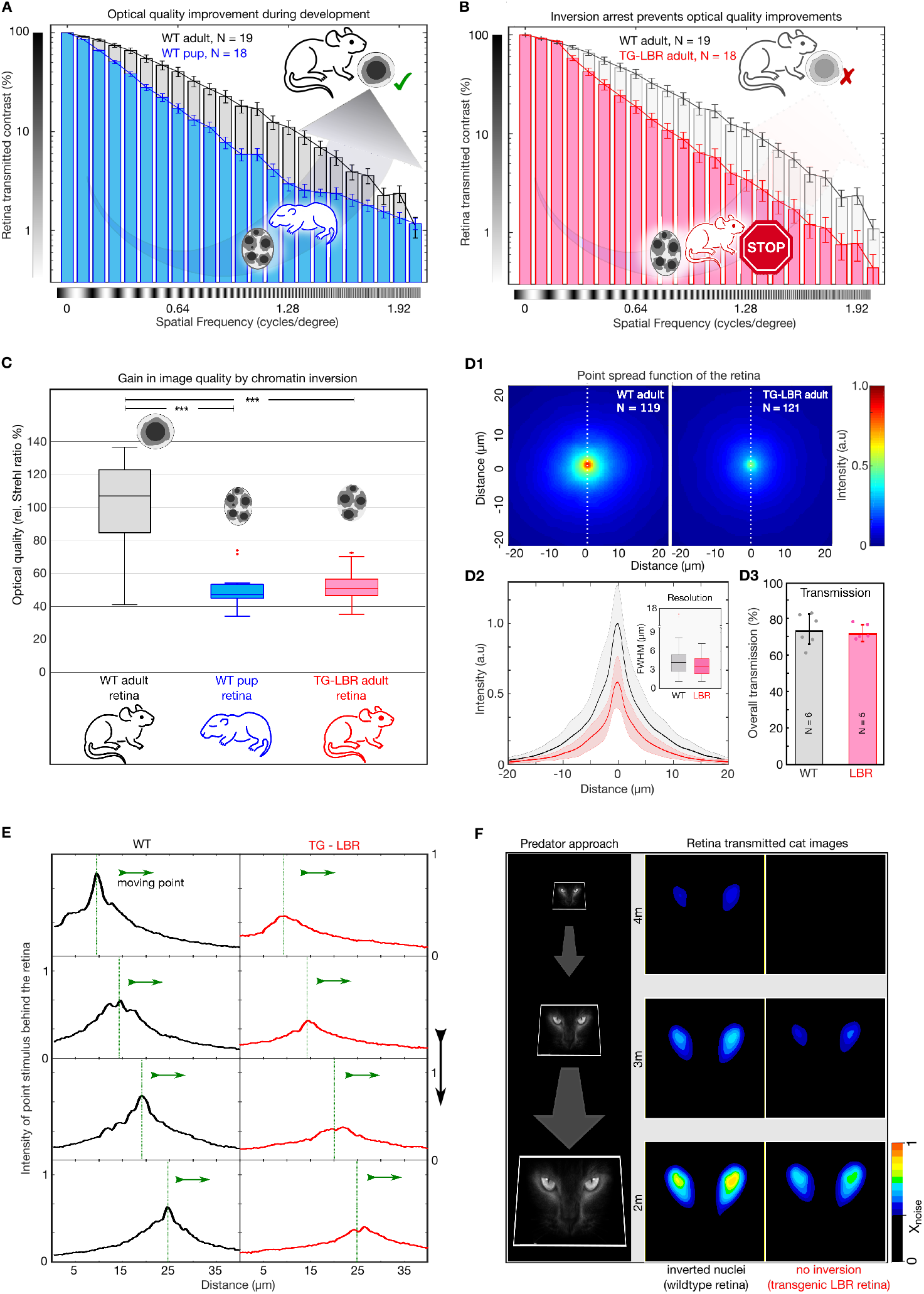
Nuclear inversion improves retinal contrast transmission characteristics. **(A)** Retinal contrast transmission increases during developmental stages of nuclear inversion, as experimentally revealed by measurements of retina-transmitted sinusoidal stripe patterns (modulation transfer functions). Developmental stage P12-14 (N=18), compared to wildtype adult (N=19 animals), note log scale. **(B)** These improvements in optical quality do not occur in retinae in which rod nuclei are transgenitically arrested in development and maintain 4-5 chromocenters. TG-LBR mouse (N=18 animals) compared to WT reference (N=19 animals), N=1950 images in total. Mean +/- 95%CI. **(C)** The optical quality improvement of the retina (relative Strehl ratios), as caused by nuclear inversion, is two-fold (p < 0.001). **(D1)** Point spread function (PSF) for WT and LBR adult retinae by projection of 3um point light stimuli through the retina, N = 240 measurements in total 6 retinae. **(D2)** Intensity quantification along the white dotted line. Shaded region shows ± 1sd. Comparable resolution in transmitted images as assessed by the FWHM of the psf **(inset). (D3)** Near identical diffuse light transmission by both WT and TG-LBR retinae (n=2 animals each, mean ± s.d.) **(E)** Intensity of a moving, retina-transmitted point stimuli for WT (black) and TG-LBR mouse (red)**. (F)** Image-series of a cat approach as seen through the retina of mice, WT and transgenic genotype from various distances at the same vision limiting (arbitrarily chosen) signal to noise level. Consistent intensity differences of two or more color shades indicate significantly better predator detection potential for WT mice. Data magnified for clarity.

Next, we asked if developmental improvements in contrast transmission of the retina are indeed caused by chromocenter fusion. For this we used mice in which LBR-overexpression largely arrested chromocenter fusion, resulting in an elevated number of chromocenters in the adult animal, similar to P14 WT (Fig. 2A2 2B), without displaying any effect on other morphological characteristics (Fig. S4). Strikingly, repeating MTF measurements on adult retinae of this inversion arrested mouse model (TG-LBR), we find near identical contrast attenuation characteristics as in the developing retina (compare Figs. 3A and 3B). Thus, developmental improvements of retinal contrast transmission are indeed mediated by the inversion of rod nuclei.

Frequently, the quality of image-forming optical systems is reported as a single parameter value called the Strehl ratio (Thibos, Hong, Bradley, & Applegate, 2004). Since our image projection setup closely mimics the mouse eye, it allows meaningful comparisons of the Strehl ratios of retinae, by comparing the volumes under MTF curves. With regard to our MTF measurements, we that find the Strehl ratio of a fully developed retina is increased 2.00 ± 0.15-fold compared to that of pups (P14) in which chromocenters fusion was not completed, and similarly 1.91 ± 0.14-fold (ratio of means ± SEM) improved compared to TG-LBR adult retinae (p<0.001) in which chromocenter fusion was deliberately arrested (Fig. 3C).

Since the Strehl ratio makes predictions for the peak intensity of a tissue transmitted point stimulus, we analyzed the effect of micro-projecting a point-like stimulus through the mouse retina (diameter here ∼3*µ*m, measurement constrained by outer segment spacing). We found that the resulting image at the back of the WT retina had a near two-fold (1.79 ± 0.38, mean ± SD) higher peak intensity compared to the TG-LBR retina (Fig. 3D, N=119, N=121, measured in at a total of 6 animals). The full-width half maximum (FWHM) of the PSF, however, remained virtually unchanged (4.32±2.38 *µ*m, 3.75±2.01 *µ*m, mean ± SD for WT & TG-LBR retinae respectively). These measurements indicate that contrast is lost due to the generation of image veil from side scattering, which overcasts attenuated, but otherwise unchanged signals. Accordingly, when comparing the integrated absolute transmission through rhodopsin-bleached retinae in dedicated experiments (Supplementary methods), we found near identical transmission values for WT and inversion arrested retinae (T_WT_ 74±8 %, T_LBR_ = 72 ±5 %, mean ±SD), which emphasizes that despite differential image signal, the overall photon arrival at the photoreceptor outer segments, remains unchanged.

The advantage of improved retinal contrast transmission becomes apparent not only when following the motion of individual (non-averaged) light stimuli that appear at considerably higher signal-to-noise levels (Fig. 3E) at the outer segments level, but also in a real-life example, when images of an approaching cat are micro-projected through a mouse retina (Fig. 3F). Nuclear inversion results in cat images becoming visible considerably earlier compared to mice that lack nuclear inversion (4 vs 3 meters, at a given arbitrary noise threshold). These results suggest that nuclear inversion may offer enhanced visual competence that originates from improved contrast preservation in retinal images.

### Improved contrast sensitivity

To determine whether the improved retinal contrast transmission translates into improved visual perception, we carried out behavioral tests using *Opto-motor-reflex* measurements (Fig. 4A). Specifically, we used a fully automated mouse tracking and data analysis pipeline (Striatech technologies) (Benkner, Mutter, Ecke, & Münch, 2013) to compare the contrast sensitivities of adult WT mice and those with arrested nuclear architecture (TG-LBR). Firstly, contrast sensitivity assessed by the animal’s ability to detect moving stripes, did not differ significantly between the two genotypes at photopic light condition (70 Lux – the typical brightness of monitor). Transgenic and WT animals showed comparable visual sensitivity, as quantified by the area under the log-contrast sensitivity curve (AULC, Fig. 4B, left)(Villegas, González, Bourdoncle, Bonnin, & Artal, 2002). As nuclear adaptation is strongly correlated with nocturnal lifestyle(Solovei et al., 2009; 2013), we adapted this set-up to assess contrast sensitivity under scotopic light conditions. At 20 mLux, which is the range of brightness in moonlight(Kyba, Mohar, & Posch, 2017), we again found comparable responses for coarse stimuli (wide large contrast stripes) suggesting equally functional rod-based vision in TG-LBR and WT mice (Fig. 4B, right) without noticeable differences in absolute sensitivity. Furthermore, mice deficient of rhodopsin (*Rho-/-*) (Humphries et al., 1997; Jaissle et al., 2001) confirmed that visual behaviour under the displayed conditions fully relies on the functionality of the rod pathway (Fig. S7E).

**Fig. 4:**
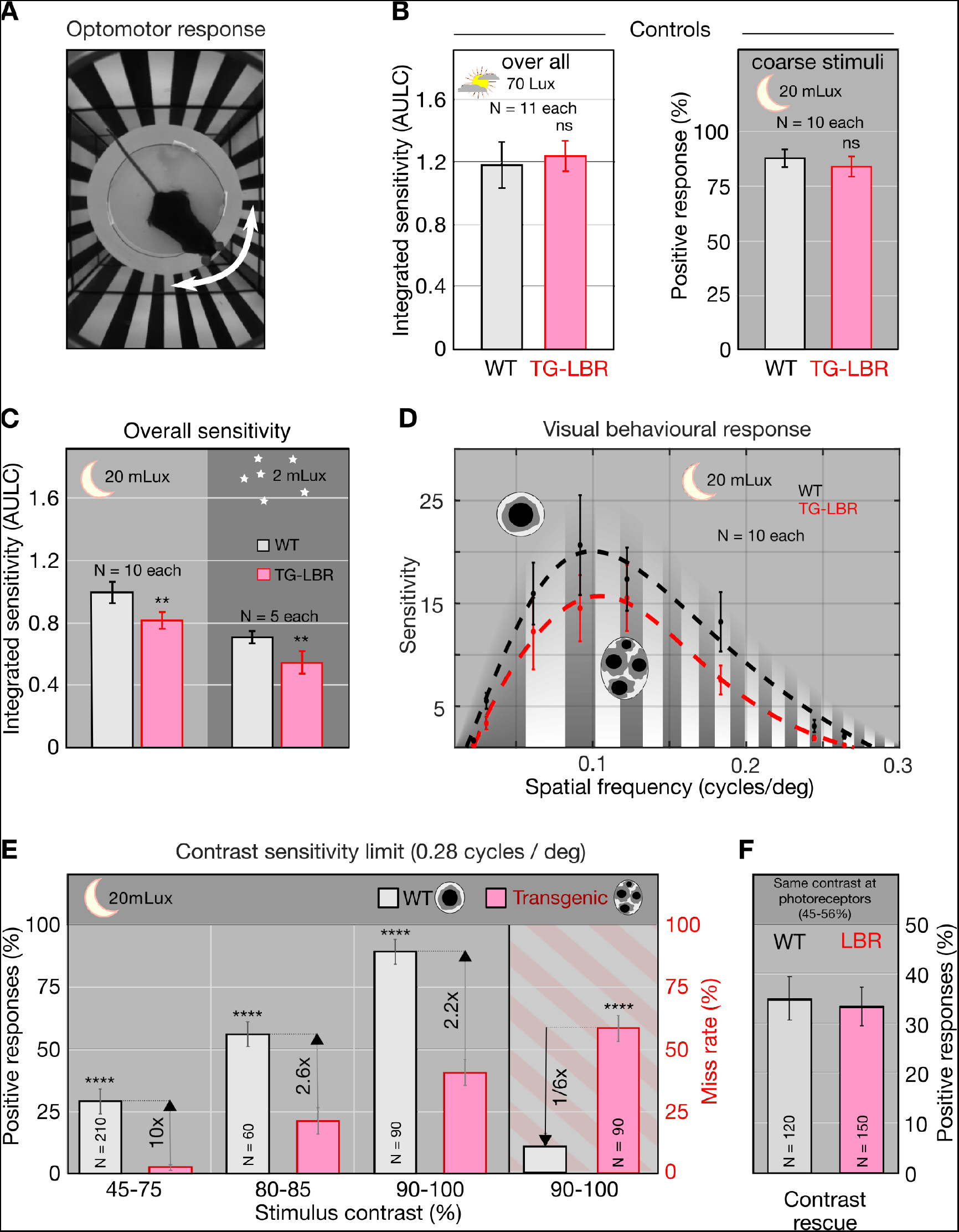
Nuclear inversion improves contrast sensitivity in the dark. **(A)** Illustration of the automated optomotor response experiment to assess the visual performance of mice, shown a 0.06 cycles/deg. **(B)** Photopic control condition and scotopic coarse stimulus (0.06cycles/deg) control showing no significant difference between WT and TG-LBR mice. **(C)** Under scotopic conditions (20 and 2 mLux) the overall sensitivity of the WT mice is 22 and 29% higher than TG-LBR mice (area under log contrast sensitivity curve, AULC). mean+/- 95% CI, (p < 0.01) **(D)** Contrast sensitivity curves evaluated at 20mLux light intensity. Significant differences appear at angular sizes above 0.15 cycles/deg. **(E)** Behaviour differences are strongest for stimuli close to the visual threshold. Here the mice in possession of the inverted rod nuclei (WT) possess an up to 10 times higher sensitivity at intermediate contrasts (29% vs 3% correct response in 45-75% contrast range), and a 6 times reduced risk to miss a motion stimulus at high contrasts (10 vs 59% failure in detection, 0.26-0.3 cycles/deg), (p<0.0001) mean +/-s.d. N indicates number of individual trials of 10 animals together for each mouse type.

When required to detect finer stripes, WT and TG-LBR mice displayed significant differences in their visual performance, specifically in contrast sensitivity (Fig. 4C). At 20mLux we observed an 18% greater AULC for WT mice compared with TG-LBR mice (p < 0.01). At even lower light intensities (2mLux, comparable to a starry night), the difference in AULC values was even greater (ratio 27%, p< 0.01) albeit at lower absolute sensitivities, which agrees with reported values for WT mice (Alam, Altimus, Douglas, Hattar, & Prusky, 2015; Prusky, Alam, Beekman, & Douglas, 2004; Prusky, West, & Douglas, 2000). The most significant differences in the contrast sensitivity occur above 0.15 cycles/degree (Fig. 4D). Especially, in the regime close to the visual acuity (0.26-0.30 cycles/degree), WT mice show up to 10 times (p<0.0001) greater positive response rates at intermediate contrasts (Fig. 4E) compared to mice with inversion arrested rod nuclei. Moreover, at 90-100% contrasts, where WT mice approach a maximum responsiveness, we observed a near 6-fold reduced risk to miss a stimulus for WT compared to TG-LBR mice (false negative rates 11% WT, 59% TG-LBR).

Finally, we asked whether reduced visual sensitivity of mice lacking the inverted nuclear architecture can be sufficiently explained by inferior contrast transmission of the retina. Direct comparison of behavioral sensitivity with the MTF curves showed that vision mostly occurs in regions in which retinal contrast transmission is higher than 50% and substantial differences in MTFs occur. Specifically, the 18-27% difference in contrast sensitivity goes together with a 26% higher Strehl ratio in WT retinae when evaluated in the relevant frequency regime (0 - 0.36 cycles / degree). This suggest that at low light level contrast sensitivity is directly limited by contrast transmission through the retina, and that a reduction of contrast sensitivity in mice with non-inverted rod nuclei may be explained by increased contrast losses in the retina. Accordingly, matching the image contrast at the photoreceptor level of TG-LBR mice with WT mice, while leaving intensities unchanged, should rescue contrast sensitivity in TG-LBR mice. To test this, we first confirmed that contrast transmission through the inner retina is a linear process, with contrasts at the photoreceptor levels being proportional to contrasts in projected images (Fig. S7 B, C). We then adjusted the displayed contrasts in optomotor measurements to pre-compensate for higher contrast losses in the TG-LBR retina. Strikingly we found that with equal image contrast at the level of the photoreceptor segments, visual competence of LBR mice was rescued and becomes near identical to that of WT mice (Fig 4F). Thus, improved retinal contrast transmission sufficiently explains the augmented contrast sensitivity in mice with inverted rod nuclei.

## Discussion

As an important determinant of fitness, animals evolved a wide range of visual adaptation to see in the dark (Nilsson, 2009; O’Carroll & Warrant, 2017; Thomas, Robison, & Johnsen, 2017; Warrant, 2017; Warrant & Nilsson, 2006). Nocturnal vision is known to rely on highly efficient light capture, both at the level of the lens and photoreceptor outer segments, and often compromises spatio-temporal resolution by summation strategies of neuronal readout (Warrant, 1999; 2017). Here we established nuclear inversion as a complementary strategy to maximize sensitivity under low light conditions. Centrally, we show that it is the direction into which light is scattered inside retinal tissue that translates into differential contrast sensitivity. Specifically, we find that the forward scattering characteristic of inverted nuclei (Solovei et al., 2009, Kreysing et al., 2010) mainly suppresses light scattering by nuclear substructure towards large angles and image veil and contrast reduction resulting from it. As the mechanism involves improvements in retinal image contrast rather than notable changes in photon transmission that could impact absolute sensitivity (Banks, Sprague, Schmoll, Parnell, & Love, 2015; Cronin et al., 2014; Nilsson, 2009; Warrant, 1999), one might ask why nuclear inversion is an adaptation exclusive to nocturnal mammals. Wouldn’t improvements in retinal image contrast not also be beneficial for diurnal mammals? Firstly, the larger spacing of photoreceptor segments in the diurnal retina significantly reduces ONL thickness and thereby the risk of scattering induced veil and loss of image contrast. Furthermore, as is well known from photography, shot-noise that accounts for image granularity (H. B. Barlow, 1956; de Vries, 1943; Rose, 1948) becomes less of a problem with increasing light levels. Million-fold higher light intensities during the day imply a higher safety margin from this noise floor (Warrant, 1999), (Fig. S7 F), as required for neural mechanisms of contrast enhancement to function (Artal et al., 2004; Flevaris & Murray, 2015; Hess & Dakin, 1988; Shevell, Holliday, & Whittle, 1992). Such compensatory mechanisms are also likely to explain why no behavioral differences are observed at elevated intensities and why augmented vision becomes pronounced only at low light levels. Last, but not least, our measurements show that, although nuclear inversion improves retinal contrast transmission via reduced image veil, resolution, the limiting factor for high acuity diurnal vision, remains largely unaffected.

In conclusion, our work adds functional significance to a prominent exception of an otherwise highly conserved pattern of nuclear organization and establishes retinal contrast transmission as a new determinant of mammalian fitness.

## Acknowledgements

We would like to acknowledge the help from the following facilities at MPI-CBG - biomedical services, transgenic core, DNA sequencing, cell technology & FACS, light-microscopy during multiple phases of the project. The authors thank Caren Norden, Jochen Guck, Elisabeth Knust, Iain Patten, Marino Zerial for helpful discussion & comments and Striatech technologies for assistance in adapting the optokinetic setup to scotopic light levels.

## Author contributions

KS and MK designed the experimental setup and experiments. IS provided the TG-LBR transgenic mouse line. MA provided Rod-Cone (Rd1/Cpfl1) double KO & Rhodopsin KO mice. KS and MK designed optical experiments, which KS carried out. KS and MK and analyzed the data. HP assisted with FACS experiments. HP, AG, KS and IS recorded morphological data. MW and EWM performed computer simulations of light propagation. KS, OB, MK, and MA designed behavioral tests, which KS and OB carried out. KS wrote initial draft. All authors contributed to a critical discussion of the data and participated in writing the manuscript, which KS and MK finalized. MK coordinated the research.

## Funding

We would like to acknowledge funding from the Max Planck Society.

## Competing interests

Authors declare no competing interests.

## Ethics Statement

“All animal studies were performed in accordance with European and German animal welfare legislation (Tierschutzgesetz), the ARVO Statement for the Use of Animals in Ophthalmic and Vision Research, and the NIH Guide for the care and use of laboratory work in strict pathogen-free conditions in the animal facilities of the Max Planck Institute of Molecular Cell Biology and Genetics, Dresden, 405 Germany and the Center for Regenerative Therapies Dresden, Germany. Protocols were approved by the Institutional Animal Welfare Officer (Tierschutzbeauftragter) and the ethics committee of the TU Dresden. Necessary licenses were obtained from the regional Ethical Commission for Animal Experimentation of Dresden, Germany (Tierversuchskommission, Landesdirektion Sachsen)”

## Supplementary Materials

### Materials & Methods

#### Retina sampling and preparation of cryo-sections

Wild type retinas were sampled from C57/BL6 mice. Eye balls of Nrl-GFP mice (Akimoto et al., 2006) were kindly provided by Jung-Woong Kim (Anand Swaroop laboratory, Ophthalmology and Visual Sciences, University of Michigan, Ann Arbor). Tissues from ROSA26-eGFP-DTA mice (Ivanova et al., 2005) were kindly provided by Dieter Saur (Klinikum rechts der Isar, Technische Universität München). Preparation of retina cryosections was performed according to protocol described earlier (Eberhart et al., 2013; Eberhart, Kimura, Leonhardt, Joffe, & Solovei, 2012; Solovei, 2010). The enucleated eye balls were shortly washed with EtOH, punched with gauss 23 needle in the equatorial plane and fixed with 4% paraformaldehyde (Carl Roth GmbH, Germany) in phosphate-buffered saline (PBS) solution for 3 hours. After fixation, samples were washed with PBS 3 x 1 h each, incubated in 10%, 20% and 30% sucrose in PBS at 4°C for 30 min in each concentration and left in 30% sucrose for overnight. The eyeballs were cut equatorially to remove the anterior parts, including cornea, lens and the vitreous, and eye cups were placed in a mold (Peel-A-Way® Disposable Embedding Molds, Polysciences Inc) filled with tissue freezing medium (Jung tissue freezing medium, Leica Microsystems). Frozen blocks were prepared by either immersion of molds with tissues in freezing medium in a 100% ethanol bath precooled to −80°C, or by placing into a container filled with precooled to −70°C 2-methylbutane. After freezing, blocks were transferred to dry ice and then stored at −80°C. Cryosections with thickness of 14–20 µm were prepared using Leica Cryostat (Leica Microsystems) and collected on SuperFrost (Super Frost Ultra Plus, Roth, Germany) or StarFrost microscopic slides (StarFrost, Kisker Biotech GmbH & Co). After cutting, sections were immediately frozen and stored in at −80 °C until use.

#### Immunostaining

Immunostaining was performed according to the protocol described in detail earlier (Eberhart et al., 2012; 2013). Prior to immunostaining, slides with cryosections were removed from −80°C freezer, allowed to thaw and dry at room temperature (RT) for 30 min and then re-hydrated in 10 mM sodium citrate buffer for 5 min. For the antigen retrieval, slides were transferred to a preheated to +80°C 10 mM sodium citrate buffer either for 5 min (H4K5ac) or for 25 min (lamin B and LBR staining). After brief rinsing in PBS at RT, slides were incubated with 0.5 % Triton X100 / PBS for 1 h, and once more rinsed in PBS before application of antibodies. Primary and secondary antibodies were diluted in blocking solution [PBS with 0.1% Triton-X100, 1% bovine serum albumin (ICN Biomedicals GmbH) and 0.1% Saponin (SERVA)]. Incubation with antibodies was performed for 12-14 hours under glass chambers in humid dark boxes (Solovei, 2010; Solovei, Grasser, & Lanctôt, 2007). Washings after incubation with antibodies were performed with PBS/0.05%Triton X-100, 3 × 30 min at 37°C. Primary antibodies included anti-lamin B (Santa Cruz, SC-6217), anti-LBR (lamin B receptor; kindly donated by Harald Herrmann, German Cancer Research Center, Heidelberg), anti-H4K5ac (kindly donated by Hiroshi Kimura, Tokyo Institute of Technology, Yokohama). Secondary antibodies were anti-mouse IgG conjugated to Alexa555 (A31570, Invitrogen) and Alexa488 (A21202, Invitrogen). Nuclei were counterstained with DAPI or Hoechst added to the secondary antibody solution. After staining, the sections were mounted under a coverslip with Vectashield (Vector Laboratories, Inc., Burlingame, CA, USA) or Aqua Poly-Mount (Polysciences, Inc., USA) antifade media and sealed with nail polish.

For microscopic analysis of FACS sorted retinal nuclei, sorted nuclei were fixed with 4% PFA in PBS for 10 mins, stained with Hoechst 33342, washed 2x with PBS and mounted on slides under coverslips in antifade medium (see below). The imaging was performed on a confocal microscopy (Zeiss LSM 700 inverted) using a Zeiss 64x 1.4 oil objective.

#### FISH

FISH on cryosections was performed as described earlier (Solovei, 2010; Solovei et al., 2007). Probes for LINE, B1 and major satellite repeat (MSR) were generated by PCR using the following primers:

5’-GCCTCAGAACTGAACAAAGA and 5’-GCTCATAATGTTGTTCCACCT for LINE1;
5’-CACGCCTGTAATCCCAGC and 5’-AGACAGGGTTTCTCTGTA for B1;
5’-GCGAGAAAACTGAAAATCAC and 5’-TCAAGTCGTCAAGTGGATG for MSR.

Probes were dissolved in hybridization mixture (50% formamide, 10% dextran sulfate, 1xSSC) at a concentration of 10-20ng/µl and hybridized to sections of mouse retina for 2 days. Post-hybridization washes included 2xSSC at +37°C (3 x 30 min) and 0.1xSSC at +61°C (10 min). Sections were counterstained with DAPI and mounted as after immunostaining (see above).

#### Microscopy and image analysis

Single optical sections or stacks of optical sections were collected using either Zeiss LSM 700 or Leica TCS SP5 confocal microscopes equipped with Plan Apo 63×/1.4 NA oil immersion objective and lasers with excitation lines 405, 488, and 561 nm. Dedicated plugins in the ImageJ program were used to compensate for axial chromatic shift between fluorochromes in confocal stacks, to create RGB stacks/images, and to arrange optical sections into galleries (Ronneberger et al., 2008)

To estimate a proportion of rods expressing LBR in retinas from TG-LBR mice, four stained cryosections from two homozygous mice were imaged. Not less than 12 image fields with pixel size of 100 nm were collected through each section. Scoring of LBR-positive and negative rods was performed in ImageJ using Cell Counter plugin. Number of chromocenters in TG-LBR rods and P14 WT pups was estimated in confocal stacks through retinas after FISH with major satellite repeat and lamin B immunostaining. Scoring of chromocenters in 210 and 65 nuclei of transgenic and P14 rods, respectively, was performed manually using ImageJ.

#### Electron Microscopy

For electron microscopy, eyes of adult WT and TG-LBR mice were fixed by cardiac perfusion with a mixture of 2% paraformaldehyde and 2.5% glutaraldehyde in 0.1M cacodilate buffer for 5 min. After eye enucleation, the eye-balls were further fixed in the same fixative for 1 h and then postfixed with OsO_4_ in cacodilate buffer for 1.5 h. Ultra-thin sections were stained with uranyl acetate and Reynolds lead citrate. Images were recorded with a megaview III camera (SIS) attached to a Philips EM 208 transmission electron microscope (FEI) operated at 70 keV.

#### Flow cytometry

FACS scattering analysis of retinal cells and sorts according to light scattering profiles were performed according to previously published (Feodorova et al., 2015). For retinae from young mice (P14 & P25) digestion times were adapted from 20 to 10 min to compensate for a faster dissociation. For brain cortical cells, digestion was preceded by vibratome slicing of freshly obtained mouse brains. N2a cells were trypsinized (0.05% Trypsin-EDTA, Thermo Fisher Scientific) and washed once with cold PBS

#### Mie models of nuclei

The scattering intensity calculations for the multi-chromocenter-nuclei depicted in Fig. 1H, were performed using Mie scattering models of spheres 0.8-4 um diameter in a refractive index contrast of 2%(Kreysing et al., 2010), the reported contrast of refraction between heterochromatin and euchromatin. Mie calculations were implemented via a modified MATLAB script (Mätzler, 2002). Relative scattering efficiencies for packed scatterers represented in Fig. S6. (E) were calculated based on dependent scattering models (Twersky, 1978).

#### Micro projection setup

Ex-vivo retinal transmission measurements were carried out using a dedicated custom built, automated optical setup. This micro-projection setup (Fig. S5. (A)) consists of two distinct optical paths, one containing projection optics (function akin to the optics of the eye) to relay images displayed by the projector LCD on to the image plane of the projection objective lens, and a second to record the retina transmitted images. The light source used (ML505L3, Thorlabs) has a spectrum close to that of the sensitivity of the rods ∼510nm. The objective lens (NA = 0.45, NPL Fluotar, Leitz, Germany) was chosen to closely match the f=-number mouse eye (Geng et al., 2011), with an additional option to narrow the incident angular spectrum for absolute transmission measurements. The projected image on the retina is then collected via an imaging / efflux objective (Olympus U PlanApo 20x 0.75/inf corr) and recorded on an Andor Zyla-5.5 sCMOS camera.

#### Calculation of MTF

To quantify Modulation Transfer Functions (MTF) we micro-projected spatially extended sinusoidal stripe patterns of different spatial frequency using a custom optical setup (Fig. S5. (A)) and recorded the transmitted images and calculate MTF from first principles. With a customized digital projector setup, the implementation of the sinusoidal stripe projection becomes straightforward analysis of wide retinal regions (Figs. S5 (A,C), Fig 6 (A-D)). The projection of spatially extended images that display many periods is however also key to capture images veil, as scattering at large angles may reduce contrast not locally (from one peak into the neighboring minimum) by across multiple periods of the test image. In industries MTF is predominantly used to assess various optical systems such as lens, cameras, displays etc. (Williams & Becklund, 1989). An advantage of MTF approach over any spatial domain approach (i.e. PSF analysis) is that overall performance of a system with optical components in series can be conveniently described as a product of the MTFs of the individual components (Boreman, 2001). In particular, MTF describes the frequency domain performance of an optical system as a ratio of the contrasts in the output image to the input object as given below,

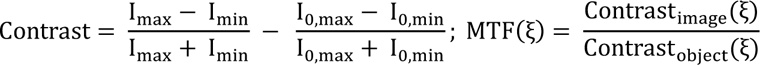

Practically, the raw images of the stripe patterns were first flat field corrected using Fiji (Schindelin et al., 2012) to ensure no global changes in contrasts affected further calculations. Each image was then processed using a custom MATLAB script by taking an average along the direction orthogonal to the contrast modulation. The resulting one-dimensional sinusoidal intensity pattern was fit to a sine wave to extract I_max_ and I_min_. (Fig. S6 A-D). Subsequently, the MTF was calculated according to the above formula. The MTF of the retina was then obtained by normalizing the measured MTF against the MTF of the optical setup alone. The differential readout of the transmitted image through the inversion arrested TG-LBR retina allows to explicitly understand the optical impact of the inner retina and the outer nuclear architecture in relation to other ocular constitutes, as lens but also photoreceptors out segments, as described in previous ex vivo (Ohzu, Enoch, & O’Hair, 1972) and in vivo studies (Geng et al., 2011; la Cera, Rodríguez, Llorente, Schaeffel, & Marcos, 2006; van Oterendorp et al., 2011),.

#### Calculation of Strehl ratio

Strehl ratio is a commonly used single number estimate of the optical performance of a system that can also be used to evaluate the optical performance of ocular components (Marsack, Thibos, & Applegate, 2004; Thibos, Hong, Bradley, & Applegate, 2003). The strehl ratio (SR) in the special domain is formally defined to be the ratio of the peak intensities of a PSF to that of a diffraction limited PSF(Strehl, 1895). In terms of the frequency domain analysis, one can more accurately calculate the SR by taking the volume under the Optical Transfer Function, albeit for systems with negligible phase transfer properties (as planar tissues), the volume under the MTF suffices to calculate the strehl ratio. This way the SR was calculated for each biologically independent sample by taking area under the frequency weighted MTF curve along the spatial frequency (Fig. S6 (F)).

#### PSF measurements

The point spread function (PSF) measurements were carried out using a 40um pinhole (P40H, Thorlabs) acting as a point light source such that, the demagnified point projected on the retina was of the size about 3*µ*m. Raw images were corrected for background by subtraction of a dark frame in FIJI. Resulting images with an ROI of 80*µ*m by 80*µ*m were normalized with respect to the integral intensity, and the central region cropped, averaged and displayed in false color.

#### Diffuse transmission measurements

The measurements were carried out with the micro projection setup above such that a point source is projected through an effective NA of 0.05 on to the retina with a final size of ∼30*µ*m diameter. The transmitted light was collected using an Olympus UPlanSApo 40x 1.25 NA silicone immersion objective lens and recorded on the camera. The fractional transmission of the samples was then calculated, after subtraction of a dark frame reference, based on the integrated intensity in the entire field of view compared to the intensity without the sample in place.

#### Hiding power

The angular-weighted integrated scattering intensity is also known as hiding power. Specifically, hiding power represents as the product of the efficiency of scattering (Q_sca_) and the directional weightage component - anisotropy factor (g) (Johnsen, 2012). The theoretical calculations based on Mie models presented in Fig.(1H, 2 F-G) were done by customizing a MATLAB script reported by (Mätzler, 2002).

#### Optical reconstitution

Equal amounts (by weight) of silica beads of diameter 2*µ*m (MSS002) and 4.5 *µ*m (MSS004a, lot obtained from Whitehouse Scientific) of RI 1.48 were dispersed in separate cuvettes containing glycerol-water mixture (RI = 1.43), the larger beads closely resembling the size of heterochromatin mass after chromocenter fusion, and the smaller beads corresponded to a ∼12 chromocenter case nuclei (at a conserved volume conserved). An edge was imaged through the two dispersions using a commercial mobile phone camera with a LED white lights acting as light source. Inset into images represent actual number ratios at a conserved volume.

#### Tissue preparation for optical characterization

Animals were scarified by cervical dislocation, and one eyeball immediately removed and opened in fresh environmentally oxygenated PBS. Next, the anterior of the eye, including the cornea and the lens, was fully removed. The retina was gently detached from the choroid, the optic nerve clipped and pulled out from the posterior cup. The retinal cup was placed on a 22×60mm coverslip. Special attention was given to remove any residual vitreous humour sticking to the retina. While the retina remained floated in PBS radial incisions were made and the retinas were flattened on the coverslip by aspiring tiny amounts of the PBS. An appropriately flattened retina was mounted under a smaller coverslip in PBS. A 255 µm spacer was placed between the two coverslips under a stereomicroscope to prevent squeezing of the retina. Preparation were typically achieved in 2 minutes, and no retina was considered for measurement with a preparation time more than 5 minutes. Optical measurements were done in an automated fashion with results in adult WT mice comparable to double pass experiments in vivo (Artal, Herreros de Tejada, Muñoz Tedó, & Green, 1998).

#### Behavioral assessment - Optomotor response

The setup to assess the visual behavioral response was obtained from Striatech technologies (Striatech UG, Tübingen, Germany) and the experiments were conducted at the CRTD. The opto-motor setup is a closed box with 4 digital displays to simulate a rotating cylinder of stripe patterns. An opening above allows the view of the animal via a camera. An independent computer-controlled software was used to track the mice on the platform. The presentation of the pattern and scoring of the tracking performance was done through a custom MATLAB program. For more details of the setup and software please refer to (Benkner et al., 2013). Age (5-6months old) and gender matched mice from Wild type and TG-LBR (transgenic) mice were used for comparison. The tests were performed under three different lighting conditions of 70Lux (Photopic), 20mLux and 2mLux (Scotopic). The visual pathway is known to be tuned for stimulus movement speed for photopic conditions and temporal variations of the contrast under scotopic conditions (Umino, Solessio, & Barlow, 2008). Accordingly, the patterns were presented at a speed of 15 deg/s for various spatial frequencies ranging from 0.01 – 0.44 cycles/deg in photopic condition. And for the scotopic conditions the stimulus was maintained at a constant temporal frequency of 0.73Hz and spatial frequency in the range 0.01 – 0.3 cycles /deg. For each contrast (100%-10% at accuracy of 5% and accuracy of 2% below 10% contrast) and size of stipe (6, 8, 11, 22, 33, 44, 55, 66, 88, 95, 100, 106 cycles/360°) the stimulus is presented for a total of 30-35s in sets of 5s each with a gap of 5s between each presentation. Once the threshold contrast was experimentally determined, for statistical analysis purpose, response values for all combinations of size and contrasts above the threshold were designated to be “yes” response and values below the threshold as “no” response. Due to limitations of the software only bar stripes were presented as opposed to pure sinusoidal stripes. Although this may affect the absolute values of threshold contrasts determined for the mice, for relative comparison of the performance of the two types of mice should remain unaffected.

#### Nocturnal adaptation of behavioural testing setup

The ambient lighting of the test chamber for photopic condition was measured using a Lux meter (Testo 540). To reduce the lighting to scotopic levels appropriate ND filter sheets (ND 3.6, ND 4.8) were placed on the monitors. The ND filters were assembled by combining ND-1.2 filter sheets (e-color+ #299, Rosco Laboratories Inc).

#### Modelling and simulation of light propagation

##### 2 -photon mapping of ONL model

Wild-type C57BL/6J mice were sacrificed by cervical dislocation. Immediately, eyes were enucleated and then cut in half around the equator, discarding all components of the eye but the eye-cup. Retina was peeled off from the eye-cup. The retinal isolation was performed in paraformaldehyde (PFA) 4% in phosphate-buffered saline (PBS) solution and then left it suspended to complete fixation for 20 minutes. The sample was transferred to a PBS solution at 4°C after fixation. The fixed sample was deposited inside a TEFLON container and embedded in low melting agarose. The agarose embedded sample was sectioned adapting the method described previously by (Clérin et al., 2014). The resulting retinal cross sections were stained with Hoechst 33342 and then wet mounted in a 50 % glycerol/PBS solution using a No. 1 cover slip (Corning). Imaging was performed with confocal microscope (LSM 780, Zeiss Germany) in two-photon mode, equipped with a tunable pulsed infrared laser (Chameleon Vision II, Coherent, US) (excitation wavelength 730nm, Objective: Zeiss LCI Plan-Neofluar 63x/1.3) resulting in an acquired intensity image of 190×190×82um with pixel-sizes of 83×83×250nm.

##### Image processing and segmentation of ONL model

To create a realistic refractive index map of packed nuclei within the ONL, we first segmented the intensity image into nuclei regions. To that end, we trained a random forest classifier via Fiji (Arganda-Carreras et al., 2017; Schindelin et al., 2012) to densely classify each pixel into background or foreground (nuclei), and applied a watershed segmentation (van der Walt et al., 2014) on the probability map with manually generated seed points, resulting in 1758 individual nuclei instances. The refractive indices for these phases have been carefully estimated previously in single cell studies (Błaszczak et al., 2014; Kreysing et al., 2010; Schürmann et al., 2017; Solovei et al., 2009). Finally, the refractive index distribution inside each nuclei region was generated according to the two different models:

Inverted: Consisting of two refractive phases with n1 = 1.357 and n2 = 1.382, corresponding to euchromatin and heterochromatin, respectively. Each nuclei mask is split into shell and core regions of equal volume (via morphological shrinking operations on each mask), which are then assigned the respective refractive indices (n1 for shell, n2 for core).

Chromocenter: Here, we randomly picked 8-12 chromocenters within the nuclei mask and assigned points close to either the nuclei border or those chromocenters to the high refractive index phase (n2) until its joint volume reached half the full nuclei volume. The other points were then assigned the less dense refractive index n1.

We blurred the resulting refractive index distribution in both cases with a small gaussian (sigma = 2px) to create a smooth distribution. For both models we furthermore ensured that both refractive phases occupied the same total volume.

##### Light propagation simulations and scattering

Light propagation through both ONL models was simulated with GPU-accelerated scalar beam propagation method (Weigert et al., 2018). A computational simulation grid of size (1024,1024,645) with pixel-size 83nm was used and the propagation of a plane wave (wavelength 500nm) through the different ONL refractive distributions was simulated. We assumed a surrounding refractive index of n_0_ =1.33. The integrated side scattering cross sections were calculated from the angular spectrum as given in (Weigert et al., 2018).

##### Relative contributions to MTFs from ONL & outer segments

In order to assess relative contribution of ONL and outer segments to the MTF of the retina respectively dedicated simulations were carried out. These compare scattering from chromocenters in to ONL (1 or 8 per nucleus), with outer segments that were simulated as 1.6 *µ*m and length 25 *µ*m cylinders (RI 1.42). For the recorded simulations the scattering anisotropy factor and efficiency was extracted, and converted into a MTF data using appropriate theoretical models (Henyey & Greenstein, 1941; Wells, 1969). Results show that outer segments only have a negligible impact on the overall MTFs (Fig (S 6G), in agreement with previous experimental findings (Enoch, 1963) and models of the OS (Vohnsen, 2007; 2014) acting as waveguides.

---

**Fig. S1.**
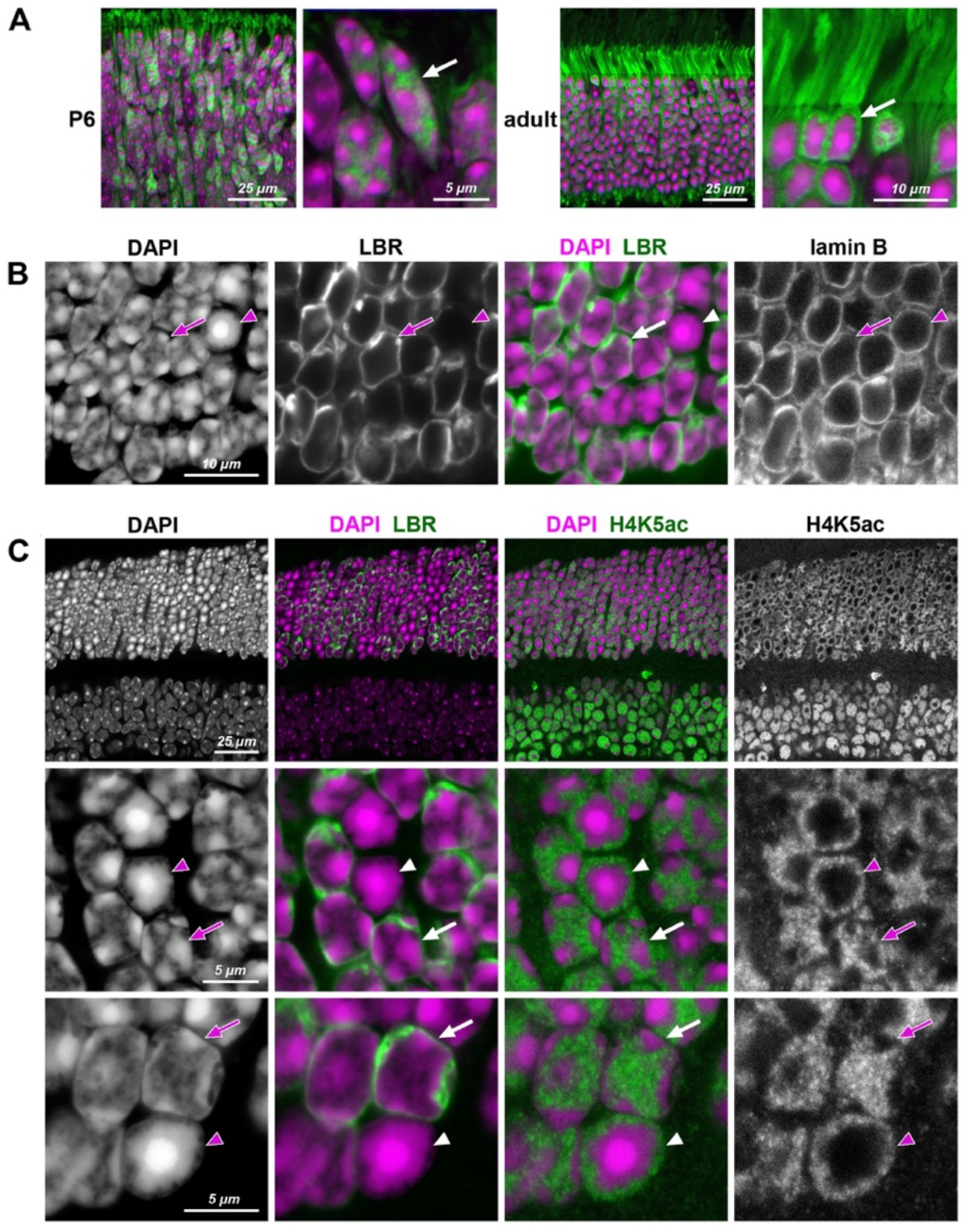
Reorganization of rod nuclear architecture in the course of postnatal retinal development (A) and in transgenic rods expressing LBR (B, C). **A**, Difference in nuclear architecture of terminally differentiated rods (adult,) and photoreceptor progenitors (P6) is highlighted by GFP (green) expressed under Nrl promoter and freely distributed through nucleoplasm and cytoplasm. During first 4-6 weeks of postnatal development, conventional nuclear architecture of rod progenitors (arrow), characterized by multiple chromocenters adjacent to the nuclear periphery, is gradually rearranged into inverted one of fully mature rods (arrow) with a single central chromocenter surrounded by LINE-rich heterochromatin. **B, C**, Rod nuclei ectopically expressing LBR (green) in adult TG-LBR retina have conventional nuclear organization with chromocenters adjacent to the nuclear lamina (B) and euchromatin occupying the nuclear interior (C). Nuclear lamina is stained with antibodies to lamin B (B) and euchromatin is highlighted by H4K5ac staining (C). Note that only proportion of rods in TG-LBR retina express LBR and thus maintain conventional nuclei (arrows). Nuclei of rods not expressing LBR are lacking peripheral tethers of heterochromatin and ultimately undergo inversion (arrowheads). Nuclei are counterstained with DAPI (magenta). Single confocal sections.

**Fig. S2.**
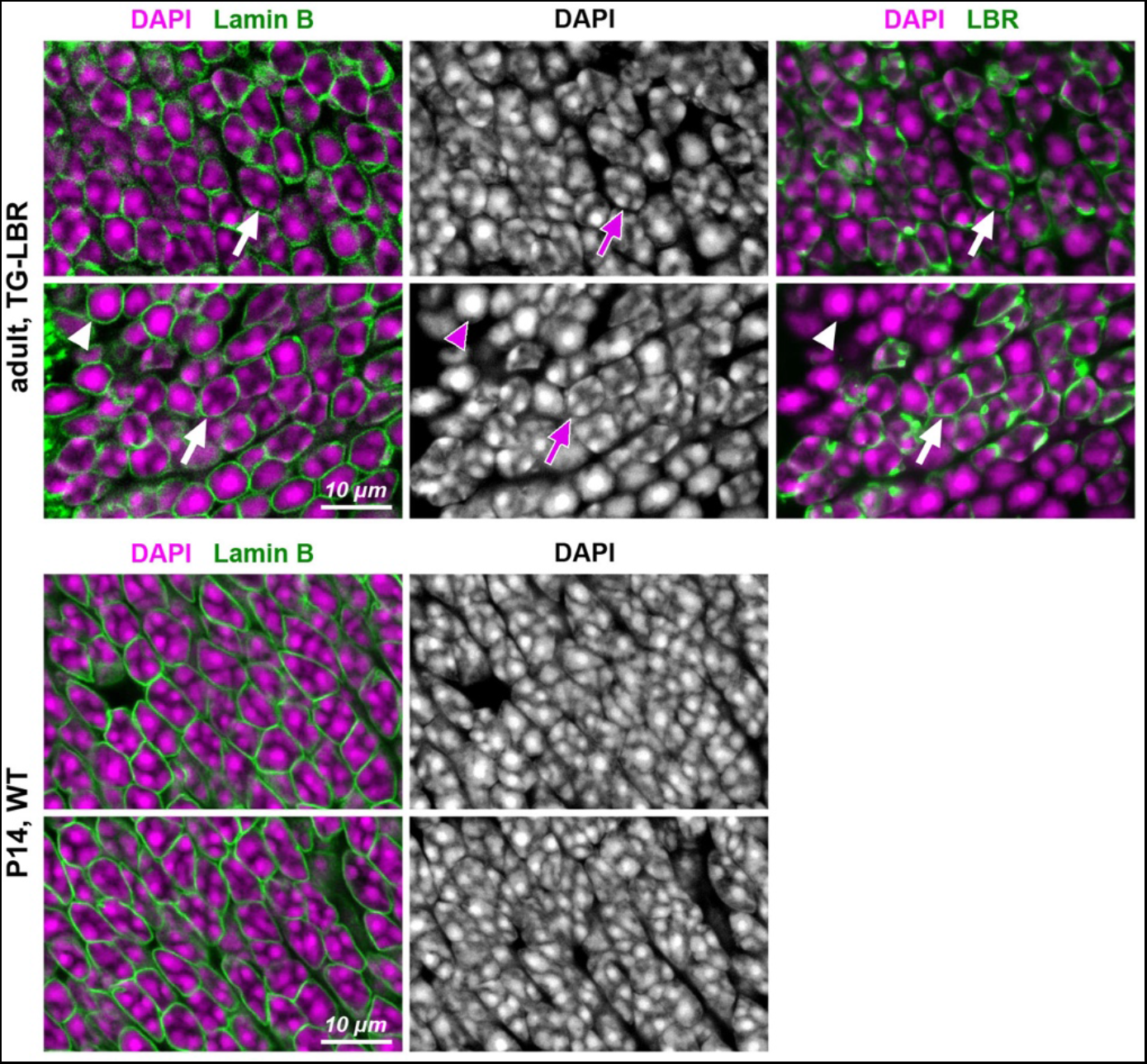
Distribution of chromocenters in nuclei of adult rods transgenitically expressing LBR in comparison to nuclei of P14 WT rods. Exemplified areas of ONL in adult TG-LBR (upper raw) and P14 WT (bottom raw) retinas. Note that only a proportion of rods in transgenic retina express LBR (see also Fig. S1). Nuclei of rods expressing LBR exhibit conventional chromatin arrangement with chromocenters adjacent to the nuclear envelope (arrows); rod nuclei not expressing LBR remain inverted with one central chromocenter (arrowheads). P14 WT rod nuclei have still conventional nuclei, although exhibit signs of ongoing inversion with massive chromocenter fusion. Immunostaining for lamin B is used to outline nuclear border (green, left panel); immunostaining for LBR is used to highlight transgenic rods (green, right panel); nuclei are counterstained with DAPI (magenta). Single confocal sections.

**Fig. S3.**
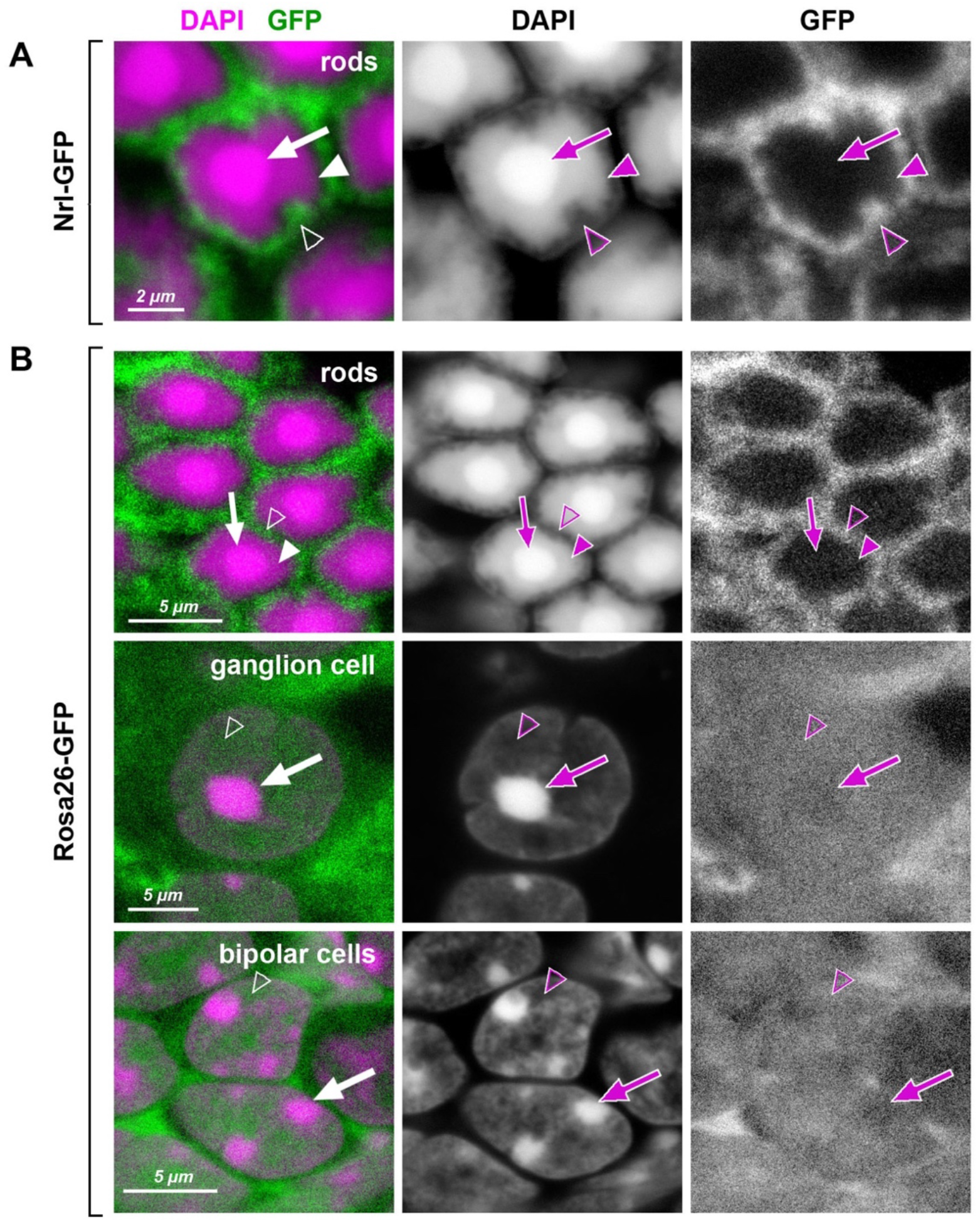
Heterochromatin in rod nuclei exhibits unusual dense packing. Retinal cells of transgenic mice expressing GFP (green) under rod-specific Nrl promoter (**A**; (Akimoto et al., 2006)) and under control of the ROSA26 promoter (**B**; (Ivanova et al., 2005)). In inverted rod nuclei, the chromatin of the central chromocenter (arrows) and the surrounding shell of LINE-rich heterochromatin (arrowheads) is packed so densely that free molecules of GFP do not penetrate into these nuclear regions. In contrast, loosely packed euchromatin in the peripheral nuclear shell (empty arrowheads) allow GFP penetration. In conventional nuclei, exemplified by ganglion and bipolar cells, the entire nucleoplasm, regardless to chromatin nature, is penetrable for GFP with chromocenters showing slightly less permeability (arrows). Nuclei are counterstained with DAPI (magenta). Single confocal sections.

**Fig. S4.**
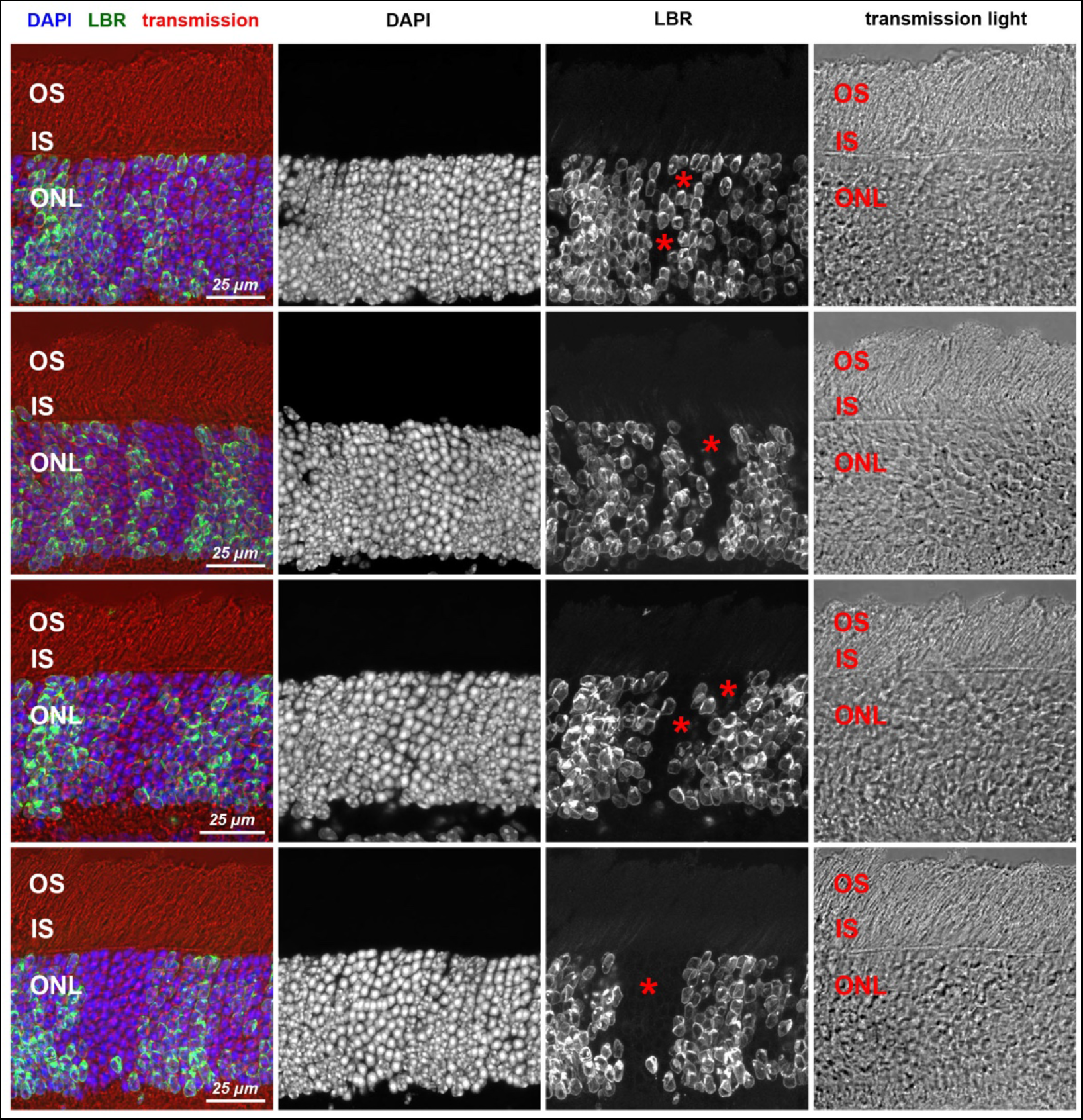
Transgenic expression of LBR does not influence rod photoreceptor structure. Retina areas with LBR-positive and LBR-negative (asterisks) rod clones demonstrating unaltered stratification. The three photoreceptor layers, represented by photoreceptor perikarya (ONL), cytoplasm of inner segments (IS) and dendritic ends of outer segment (OS), are preserved in TG-LBR retina. Immunostaining of LBR (green); nuclear counterstain with DAPI (blue). Maximum intensity projections of 5-7 µm confocal stacks.

**Fig. S5.**
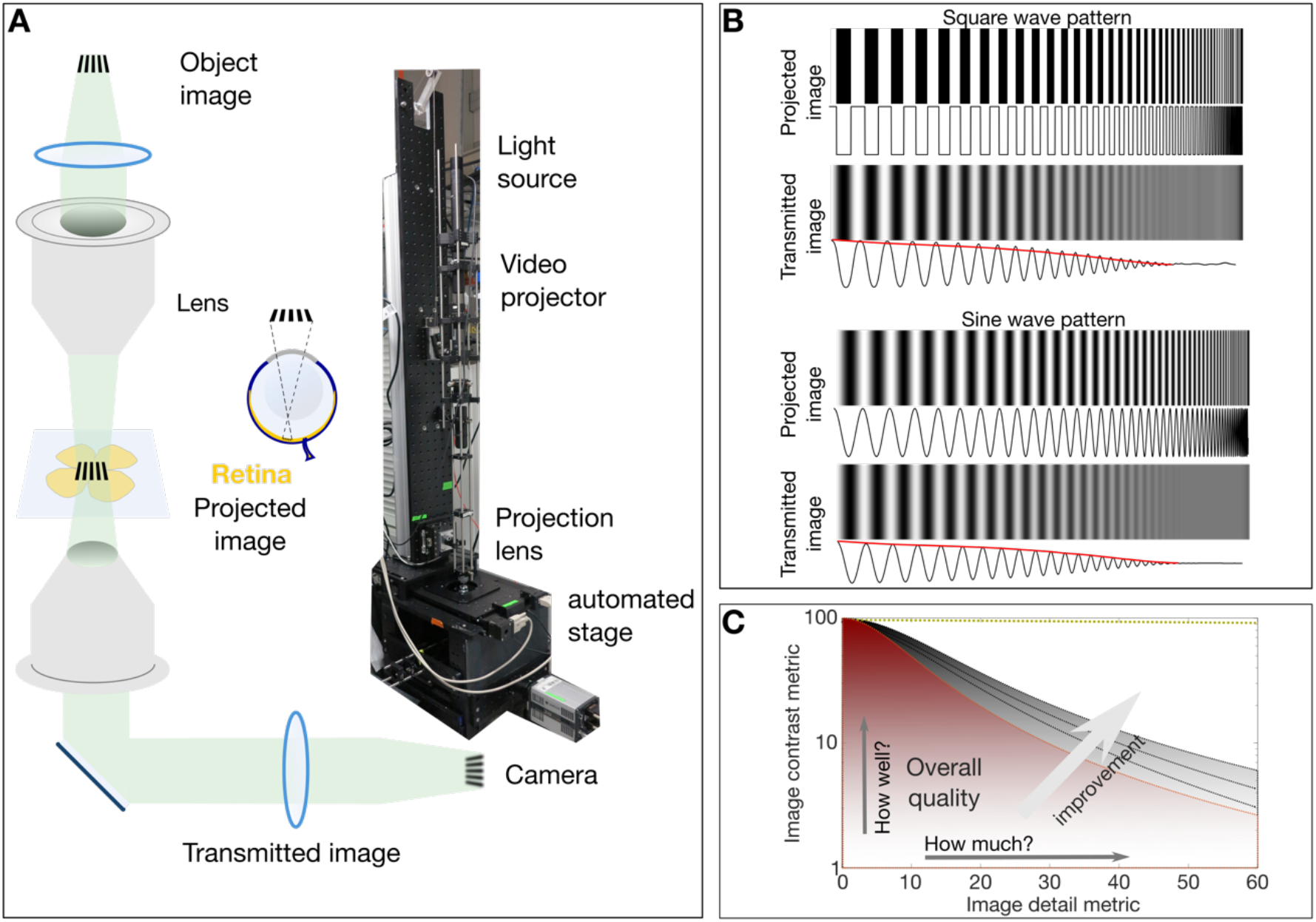
Simplified schematic of the custom micro-projection setup and the concept of modulation transfer function. **(A)** Simplified schematic and photo of the micro projection setup indicating the image object. The projecting lens functions like the biological eye (same NA = 0.45) to project the image on to the retina. The transmitted image is collected by a second lens and recorded on a camera. **(B)** The loss of contrast inherent to imaging systems (compare contrasts and intensity signals in projected and transmitted images). The envelope of the gradually degreasing signal (shown in red) is essentially the transfer function of the imaging system. The study of the contrast at various image details and fineness results in the MTF curves as shown in (c). **(C)** A typical MTF plot of image contrast vs image detail parameters. In this depicted case the overall image quality is proportional to the area under the curve that can yield the Strehl Ratio (a combined metric of how well and how much of the image is visible).

**Fig. S6.**
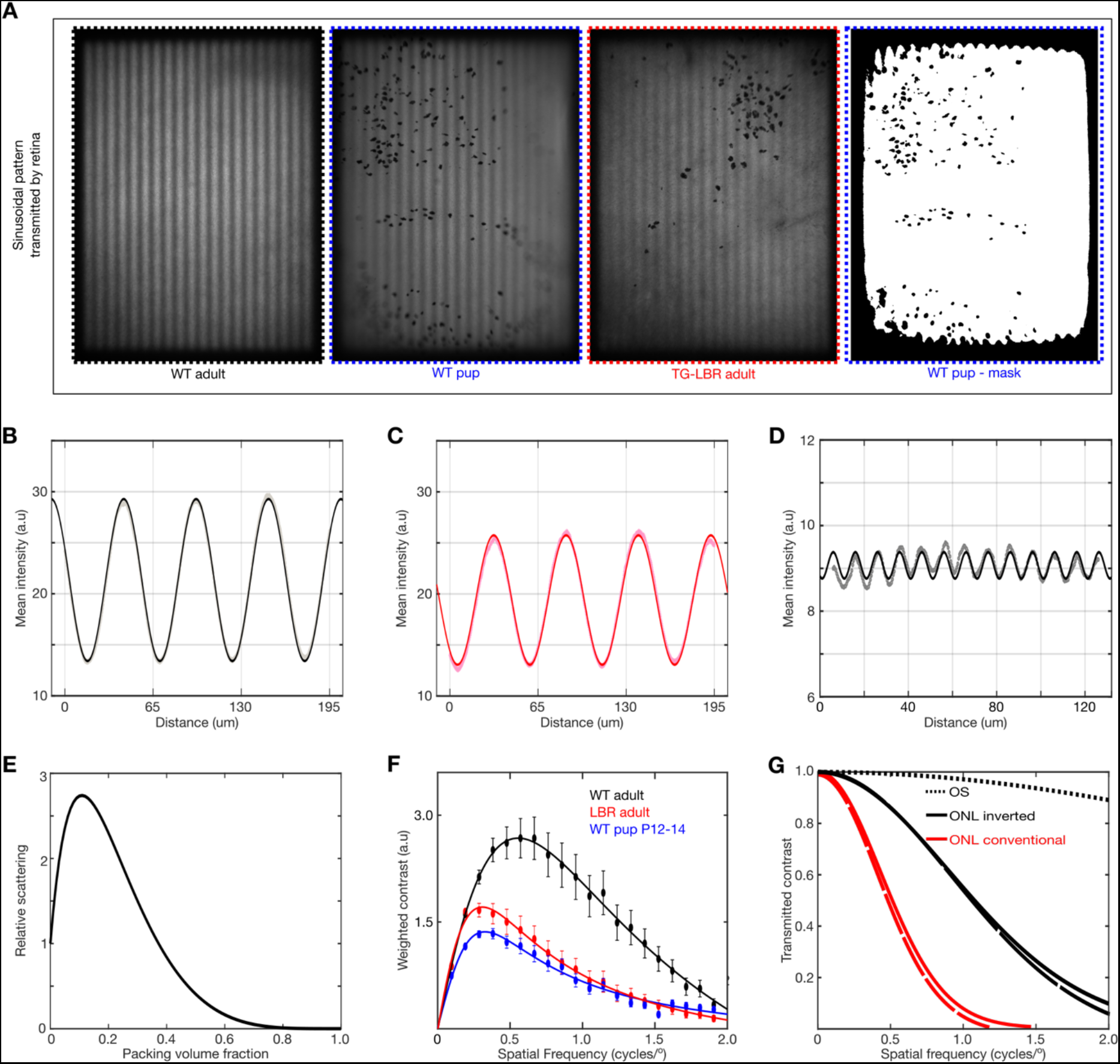
Modulation Transfer Function and its relation to light scattering and visual perception. **(A)** Representative images of sinusoidal patterns transmitted by the different retinae and a mask image (far right) illustrating the ROI used for contrast analysis. **(B, C)** Fitting of pure sinusoidal curves to the measured intensity in a WT and TG-LBR mouse retina at 0.1cyc/deg. The greater transmission of contrast is evident from the amplitude of the sine waves in the WT retina. **(D)** Illustration of robustness of the sine curve fitting for very low residual contrast (∼3%) and very fine image details. **(E)** Dependent scattering effects due to close packing of scatterers for various volume fractions. (Calculations based on models described in literature). **(F)** Frequency weighted contrast transmission curves for the evaluation of the Strehl Ratio. **(G)** Estimates of comparison of MTF from modelling and simulation of light scattering by ONL and OS illustrating the dominant effect of the packed ONL as opposed to the OS.

**Fig. S7.**
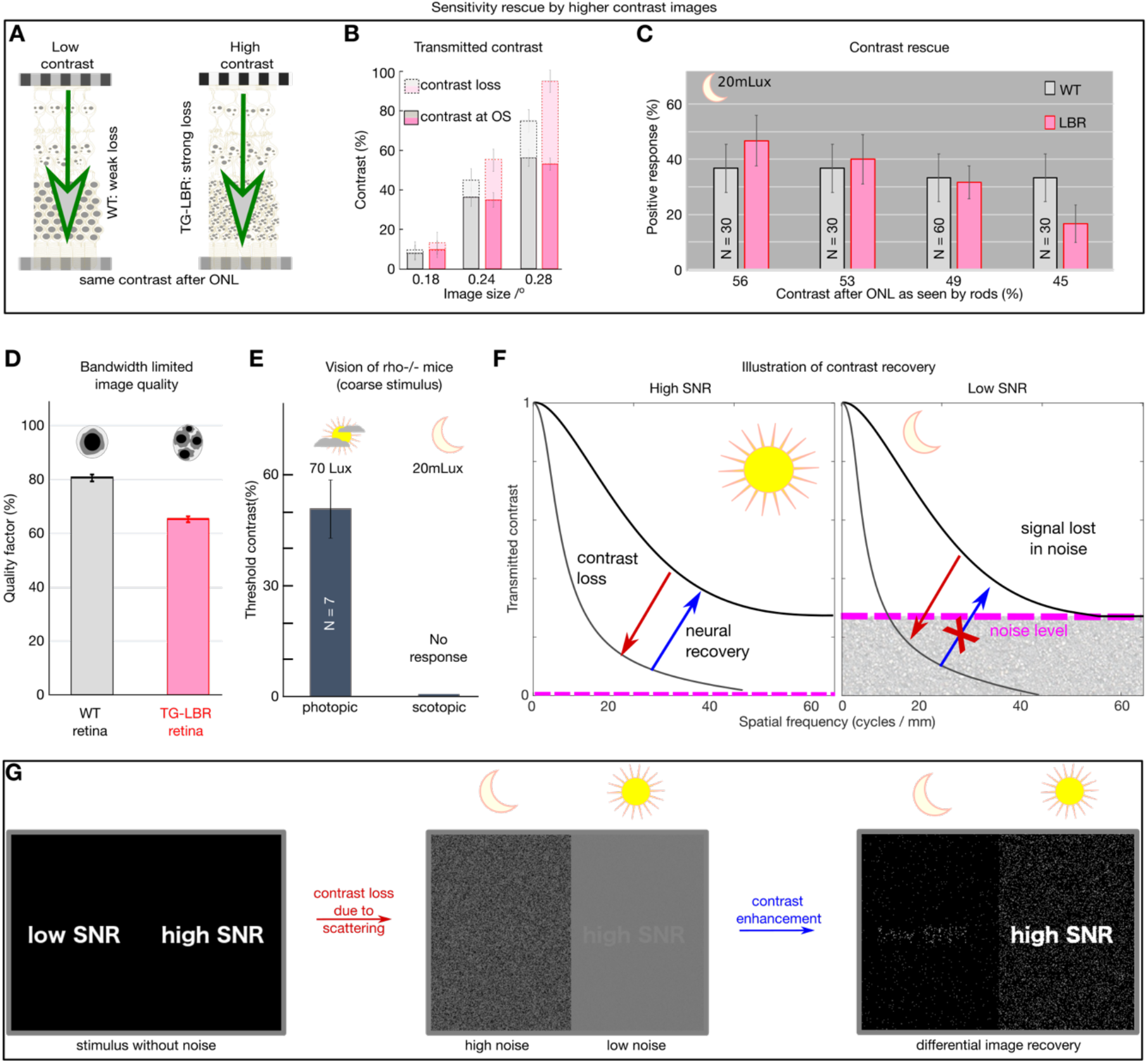
Retina transmitted contrast directly impacts visual behaviour. **(A)** Illustration of the role of retina as an information filter and contrast loss buckets for WT and TG-LBR retina. **(B)** Stimuli contrast for OMR response presented such that after corresponding loss in the WT and TG-LBR retinae, the OS receives comparable contrast of ∼9, 36 and 54% at 0.18, 0.24, 0.28 cycles/deg respectively for transduction. **(C)** The visual response of the TG-LBR mouse is recovered to the level of WT mouse by a stimulus with higher input image contrast compensating for the greater loss due to its ONL with highly scattering non-inverted nuclei. **(D)** Estimates of image quality for the corresponding mice based on the MTF of WT and TG-LBR retina at relevant range of visually sensitive image details (0.15-0.36 cycles/degree). **(E)** Control to verify the scotopic illumination settings of the OKT setup, depicting no response for a rhodopsin KO mouse. **(F)** Day and night vision differ significantly in the signal to noise level (Warrant, 1999), and loss of signal below the noise floor is lost from an information point of view and cannot be recovered. **(G)** Illustration of the contrast recovery for stimulus under low and high noise conditions. The scattering induced contrast loss and image degradation in a noisy low light environment cannot be recovered by any contrast enhancement mechanism

**Fig. S8.**
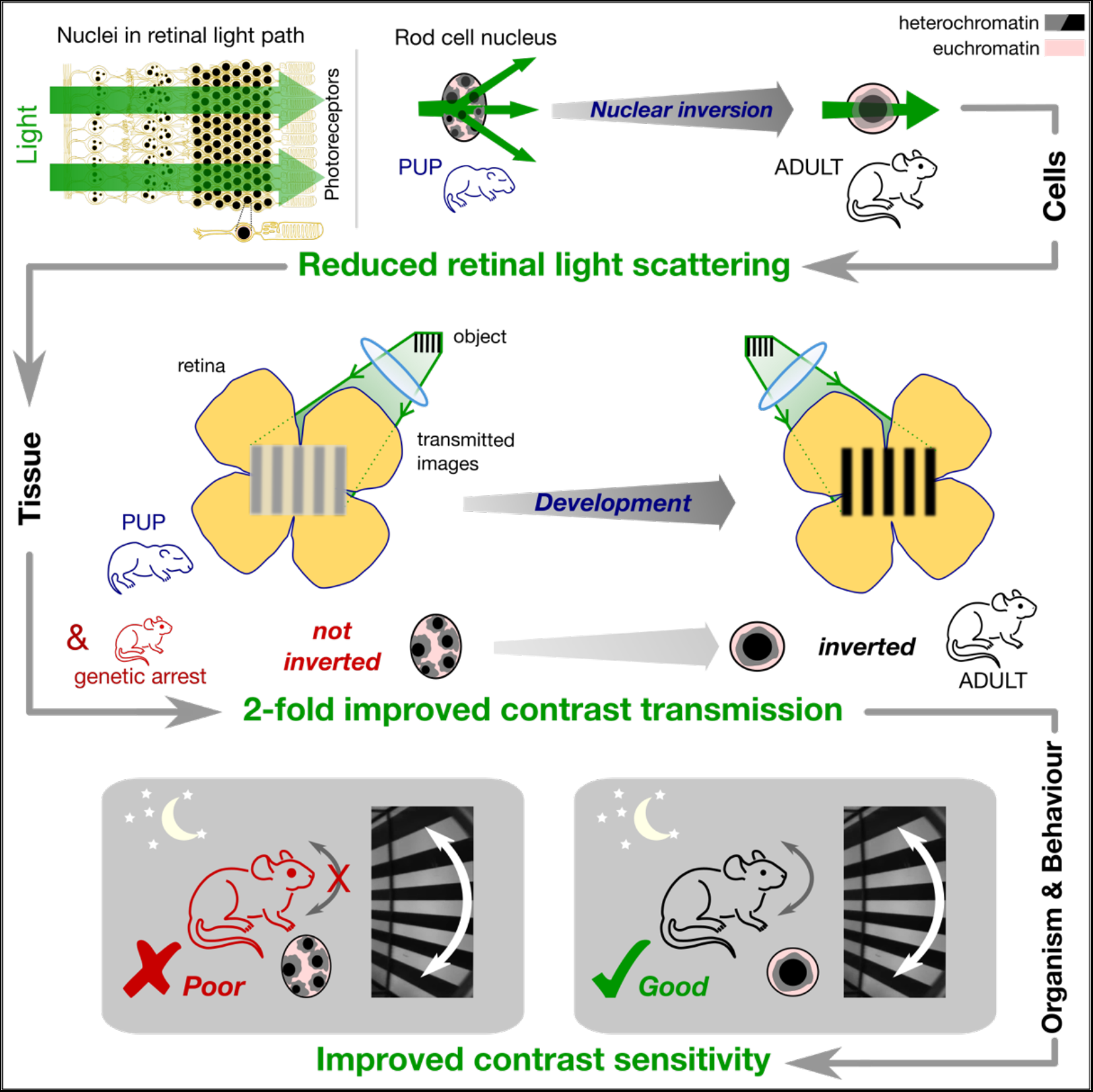
Model of nuclear adaptation enhancing nocturnal vision. The abundance of rod nuclei in the nocturnal mammalian retina presents a significant barrier to light. The fusion of heterochromatic chromocenters reduces the volume specific scattering mainly at large angles. This results in a reduced light scattering at the tissue level leading lower scattering induced veil. Reduction in light scattering leads to a near two-fold improvement in the contrast transmission by the retina. Improved image quality transmitted by the retina finally enables greater contrast sensitivity exclusively at nocturnal conditions.

**Movie S1.**

2 photon volumetric image of WT mouse retina.

**Movie S2.**

3D morphological models of ONL RI distribution used in light propagation simulations.

**Movie S3.**

Behaving mouse in an Optomotor response set up.

## Notes

#### Summary of Updates

Manuscript file pdf with Supplemental files updated

